# Visual self-motion feedback affects the sense of self in virtual reality

**DOI:** 10.1101/636944

**Authors:** Aubrieann Schettler, Ian Holstead, John Turri, Michael Barnett-Cowan

## Abstract

We assessed how self-motion affects the visual representation of the self. We constructed a novel virtual reality experiment that systematically varied an avatar’s motion and also biological sex. Participants were presented with pairs of avatars that visually represented the participant (“self avatar”), or another person (“opposite avatar”). Avatar motion either corresponded with the participant’s motion, or was decoupled from the participant’s motion. The results show that participants identified with i) “self avatars” over “opposite avatars”, ii) avatars moving congruently with self-motion over incongruent motion, and importantly iii) identification with the “opposite avatar” over the “self avatar” when the opposite avatar’s motion was congruent with self-motion. Our results suggest that both self-motion and biological sex are relevant to the body schema and body image and that congruent bottom-up visual feedback of self-motion is particularly important for the sense of self and capable of overriding top-down self-identification factors such as biological sex.

## Introduction

Humans have the conscious experience of having a sense of self (Gallagher, 2000). In the last twenty years, there has been a rise in experimental research on bodily self-consciousness and the sense of self (Blanke et al., 2015). At the same time there has been a rise in the use of virtual reality (VR), which has enabled researchers to manipulate visual feedback of the self, using virtual avatars in order to assess vision’s role on the body schema (Maravita and Iriki, 2004) as well as body image (Mölbert et al., 2018). Although there is no consensus among researchers of these terms, body schema is generally regarded as an unconscious, bottom-up, dynamic representation, relying on proprioceptive information from the muscles, joints, and skin during self-motion (Gallagher, 2000). On the other hand, the body image is a more conscious, top down, cognitive representation, incorporating semantic knowledge of the body and one’s identity including biological sex (Longo et al., 2008). An interesting research question arises then: to what extent does visual self-motion feedback and biological sex identification affect the sense of self in virtual reality?

Previous experiments such as the rubber hand illusion (Tsakiris and Haggard, 2005; Ehrsson et al., 2005; Suzuki, K et al., 2013) and whole-body illusions (Aglioti and Candidi, 2011; Petkova and Ehrsson, 2008; Blanke and Metzinger, 2009) have demonstrated that vision can powerfully change the representation of the self where ownership can even be induced over a body of a different race, age, or gender (Petkova and Ehrsson, 2008; Maister et al., 2014). Here, it is the temporal coupling of synchronous touch and vision of being touched that enables one to feel embodiment and a sense of agency over another body or body part. Similarly information about self-motion from proprioception and the vestibular system has also been shown to contribute to high-level bodily perception, such as the sense of body ownership (Lopez, 2015). It has also been suggested that the interaction between vestibular and bodily somatosensations is a form of self-representation, which is important to link the spatial description of one’s own body to the spatial description of the outside world (Ferrè & Haggard, 2015). Further, out-of-body experiences, in which patients localise the self outside their own body and experience seeing their body from a disembodied location, have been attributed to failures in integrating multisensory bodily information due to conflicting visual and vestibular information (Kaliuzhna et al., 2015). Finally, while experiments have recently shown that bodily self-consciousness may arise from the integration of multisensory signals in fronto-parietal and temporo-parietal regions of the brain (Blanke et al. 2015), much more work remains to be done to determine the plasticity of human bodily self-consciousness.

Self-motion is critical to how we identify sensory experiences related to the body with our identity. Indeed self-recognition among humans, as well as and other species (Gallup, 1970; Reiss and Marino, 2001; Ari and D’Agostino, 2016) and perhaps even machines (Hart and Scassellati, 2012), is conventionally identified by the understanding that actively exploring one’s own mirror reflection enables us to recognize that the reflected image of oneself does not represent another individual but oneself. Like a mirror, VR’s interactive computer interface that immerses the user in a synthetic 3-dimensional environment can give the user the illusion of identifying with the visual representation of the body and “being” in that virtual setting through a sense of “presence” (Sanchez-Vives & Slater, 2005). VR technology is uniquely capable of being used to assess human self-awareness by presenting the user with an avatar representation of their self. Here resemblance of the artificial body to a human body improves embodiment into the avatar (Maselli and Slater, 2013), and the feeling of presence in the virtual world (Eastin, 2006).

Here we sought to assess the role of self-motion and the sense of self by presenting participants with pairs of mirrored avatars that visually represent their self, or another person. Self-motion was manipulated by assigning one of the avatars to move 1:1 with participant movement using motion tracking, or alternatively by adding different motions to participant movement to present realistic but decoupled motion. Through various combinations of avatar pairs and motion applied to each avatar, we assess the hypothesis that participants’ judgements of their sense of self will be influenced by both visual self-motion and biological sex of encountered avatars.

## Materials & Methods

### Participants

Nine participants (7 males; 24-54 years; average age: 27.2 years; average height: 173.5cm SD = 5.09cm) participated in the experiment. All participants were Korean and visiting the Games Institute at the University of Waterloo as part of a workshop series on gaming. All consent materials and instructions were translated into Korean. A Korean translator familiar with the experiment was available at all times to ensure that participants understood experimental instructions. Participants had normal/corrected to normal vision and had no known neurological disorders. The experiment was approved by the University of Waterloo’s Research Ethics Committee, which complies with the Declaration of Helsinki.

### Avatar creation

As recent research suggests that the customization of avatars increases the extent to which people feel connected to their avatars (Lim and Reeves, 2009), participants in our experiment created their own avatars. In the first stage of the experiment, participants had thirty minutes to create a computer-generated avatar that they felt most represented their self using Adobe Fuse Creative Cloud (https://helpx.adobe.com/creative-cloud/how-to/create-3d-character-adobe-fuse.html; Figure 1). This software enabled participants to rapidly assemble a 3D character from more than 20 base characters and further customize it into a unique character with different weight, height, skin tones, and texture to their satisfaction. A mirror was provided for each participant for reference and experimenters as well as the translator were available to technically assist participants when needed. All participants reported that they were satisfied with their avatars within the thirty-minute creation period. After participants created their “self” avatars, the experimenters created “other” avatars that were of the opposite biological sex of each participant by swapping each body segment while closely matching size. The clothing used on the other-avatar was matched as close as possible to the clothing on the self-avatar, but some were more different than others depending on the initial clothing selected by each participant (Figure 2).

**Figure 1:**
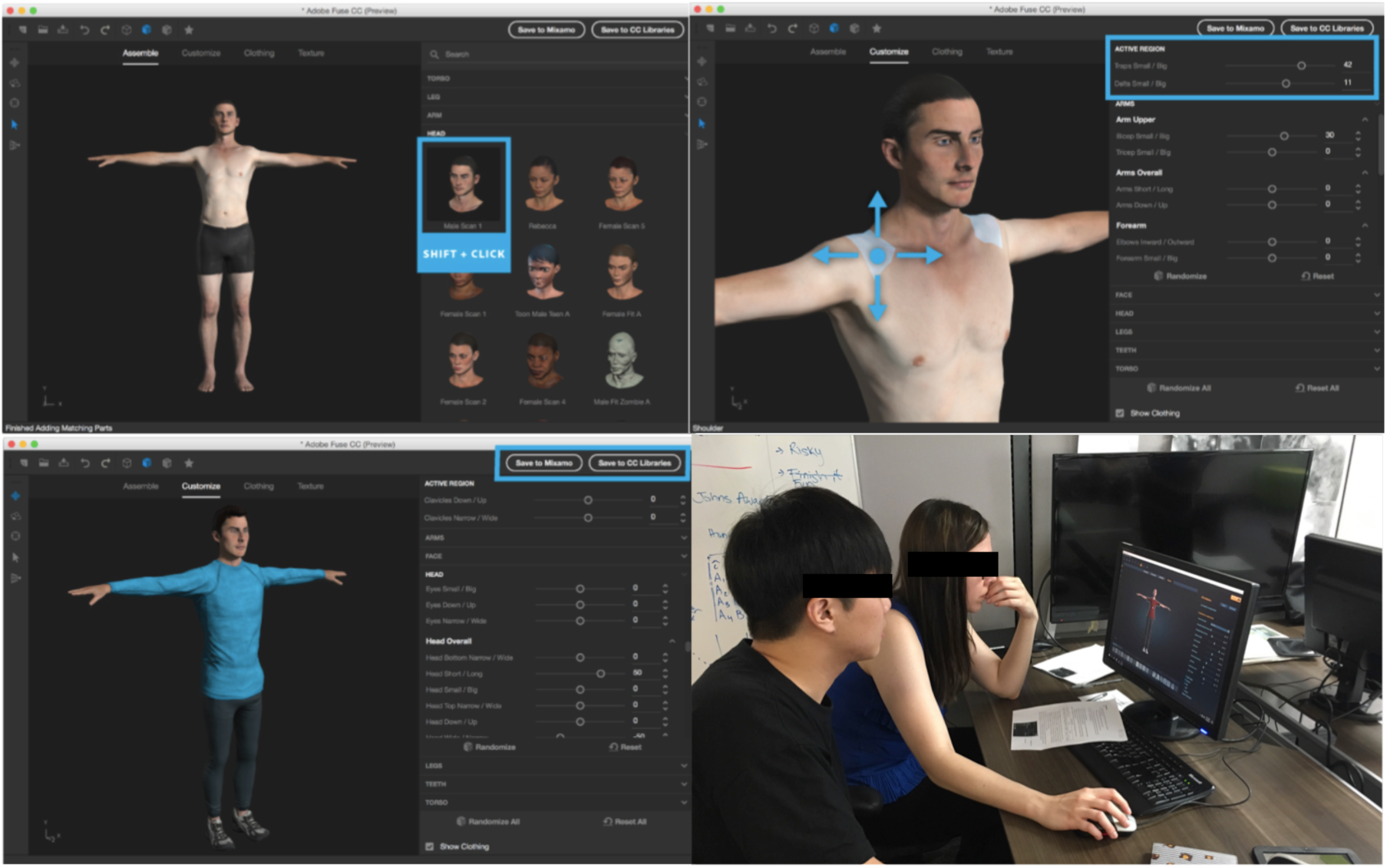
Participants had thirty minutes to work with Adobe Fuse software to create an avatar that they felt most closely represented their self. Participants were provided a mirror for reference. Following completion, the experimenter swapped all body components with those of the opposite sex of the participant in consultation with the translator but not the participants themselves (lower right). Written informed consent was obtained from the individuals for the publication of this image. Note however that these individuals have faces obscured for this preprint.

**Figure 2:**
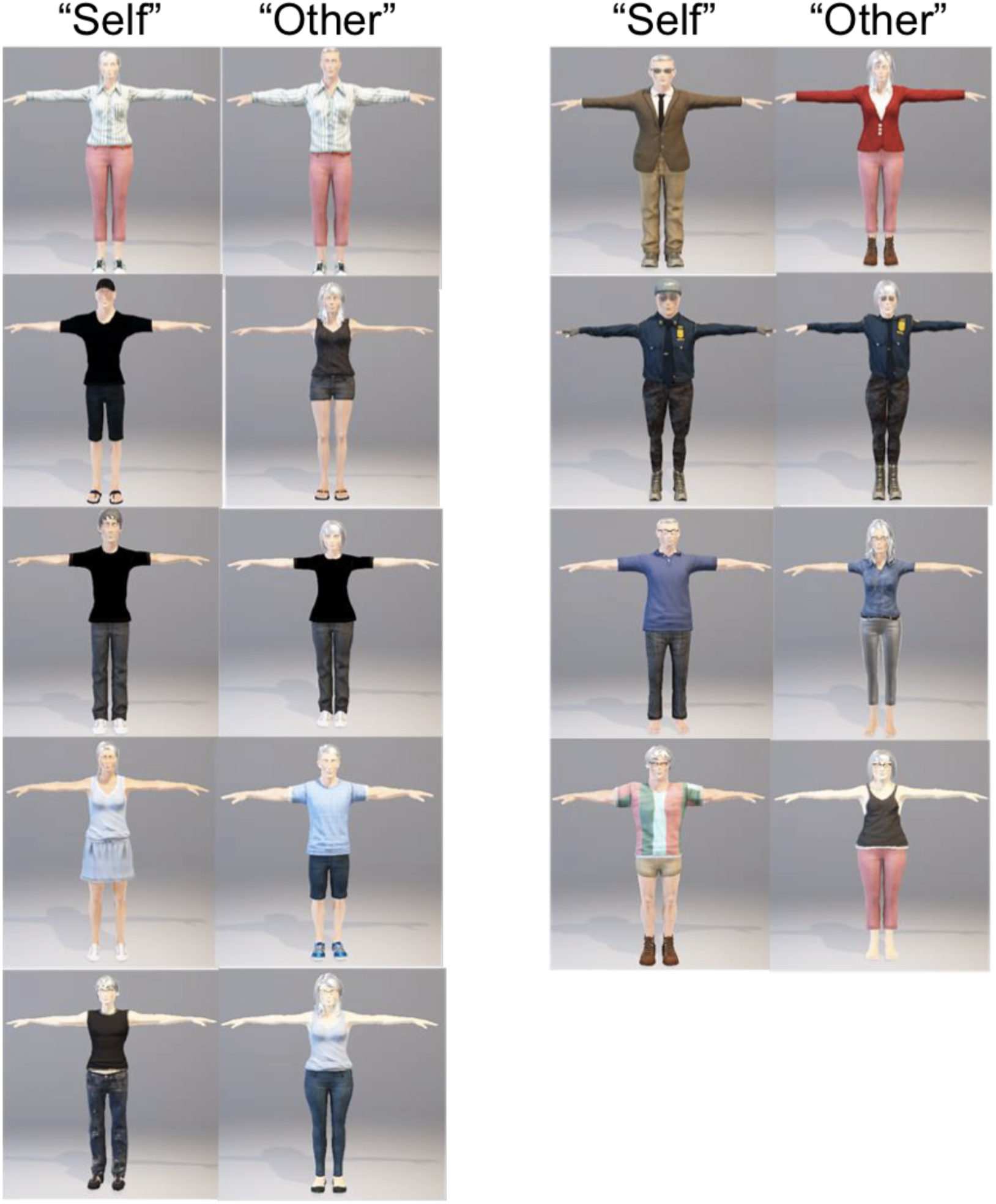
Avatars created by each of the nine participants (“self”), and avatars that were created in the image of the opposite sex (“other”) by the experimenter.

### Equipment

Participants were fitted with the HTC VIVE virtual reality head mounted display (VRHMD; Figure 3). The HTC Vive uses an organic light-emitting diode display and provides a combined resolution of 2160×1200 (1080×1200 per eye) with a refresh rate of 90 Hz and a field of view of about 110 degrees. The VRHMD weighs about 555 grams and was connected by link box that supplied power, HDMI 1.4, and USB connection to a PC that generated an environment that was rendered by a high-end graphics card (NVIDIA GTX1070). Two stationary reference cameras were used to track the participant’s head while they moved within the virtual environment (Figure 4; https://www.htcvive.com). The Microsoft Kinect One (v2) motion capture camera (https://developer.microsoft.com/en-us/windows/kinect) was used to capture movement of the participants’ whole-body motion. The Kinect is a motion sensing device used for video game interaction and computer interfacing that performs body pose tracking based on an infrared time of flight camera. Its SDK contains drivers and an API, which allows accurate body pose inference, in the form of an animated skeleton, and scripts exemplifying application development that enabled us to animate the avatars used in the experiment. The VRHMD was equipped with a Leap Motion controller (Figure 4: https://www.leapmotion.com) mounted to the front to capture high-resolution hand and finger movement tracking. The Leap Motion controller is a small (7.5 × 2.5 × 1.2 cm) sensor using 3 high-powered infrared LEDs and two high-resolution cameras for hand tracking. Its SDK uses undisclosed algorithms based on stereo images to track two hand skeletal models that we integrated with body tracking from the Kinect.

**Figure 3:**
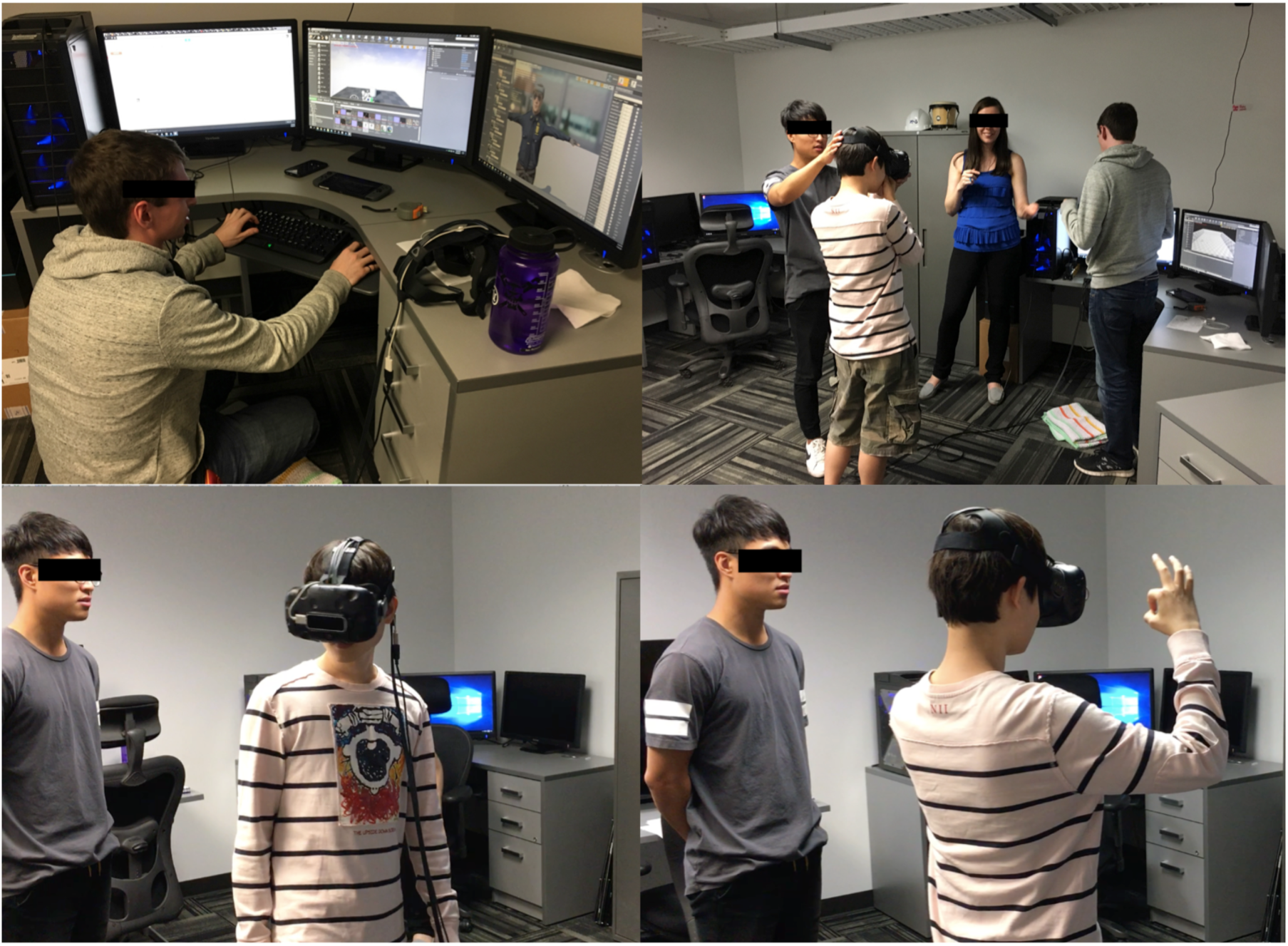
Avatars created in Adobe Fuse were imported into the virtual environment using the Unreal Engine 4 by the experimenter (upper left). Participants were equipped with the HTC VIVE head mounted display (upper right), which was equipped with the Leap Motion camera (lower left). Participants were given instructions in Korean by a translator and trained to indicate whether they preferred the avatar on the left or right by raising their corresponding left or right hand (lower right). Written informed consent was obtained from the individuals for the publication of this image. Note however that these individuals have faces obscured for this preprint.

**Figure 4:**
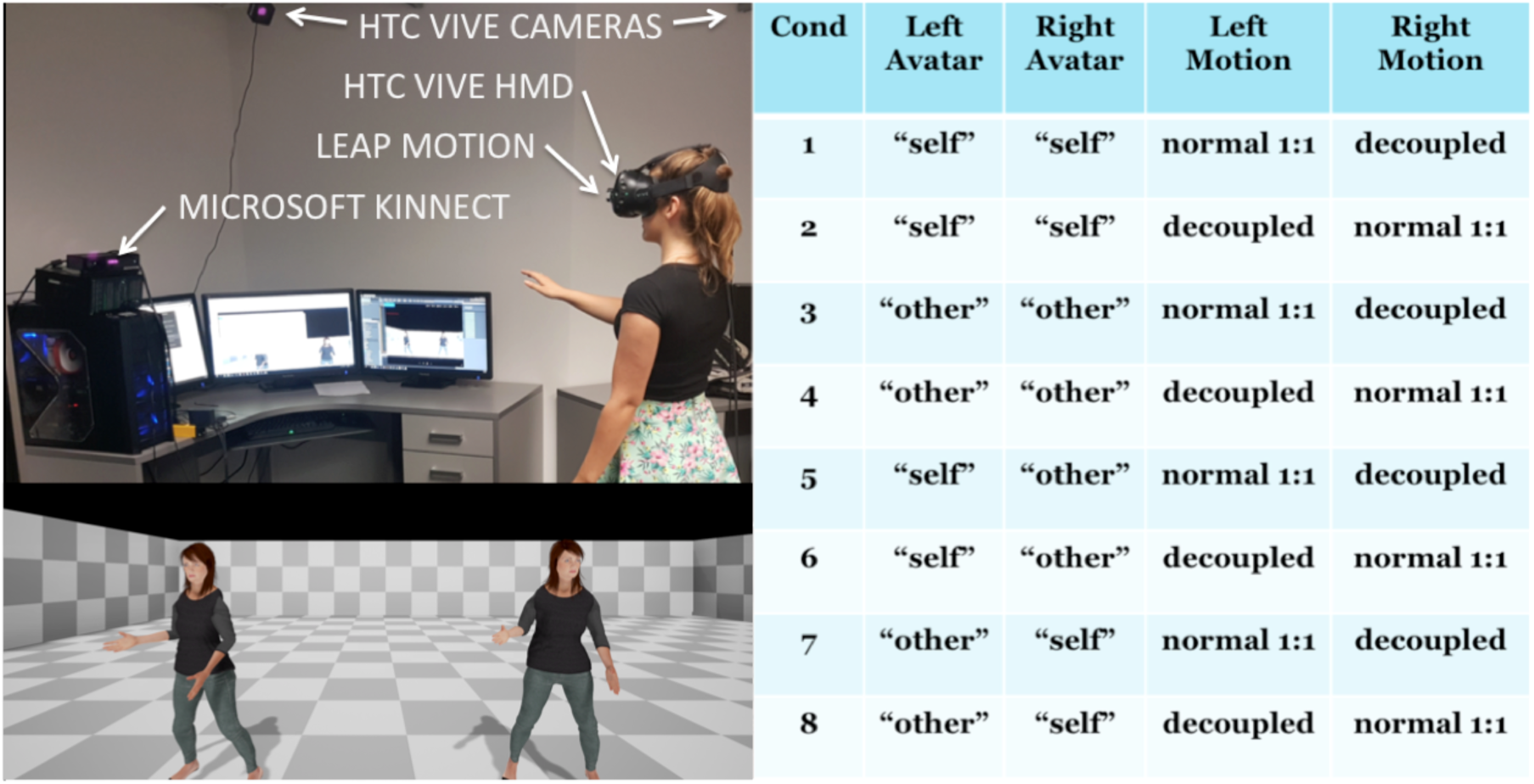
Experimental equipment (upper left) and design (right). Normal 1:1 self-motion was captured through combination of the HTC VIVE inertial measurement units, HTC VIVE cameras, Leap Motion camera, and Microsoft Kinnect camera. On each trial, participants were presented with two avatars (lower left) rendered two virtual meters in front of them, one of which always had normal 1:1 motion and the other with “decoupled motion”. Decoupled motion was generated by averaging normal 1:1 motion with an “idle pose” avatar available from Mixamo and imported into the Unreal Engine 4 virtual environment. Written informed consent was obtained from the individual for the publication of this image.

A game engine (Unreal Engine 4, https://www.unrealengine.com) was leveraged for its built-in rendering, simulation, and networking capabilities, while the processing of inputs listed above that captured self-motion were accomplished by using plugins to the game engine. “Self” and “other” avatars created with Adobe Fuse were transferred into Mixamo software (https://mixamo.com) which enabled animation of avatars (Figure 4) while also allowing us to subtly decouple self-motion from true 1:1 motion by averaging true 1:1 motion with subtle animation of a “breathing idle” animated character.

### Task

On each trial, participants were exposed to a simple virtual reality scene with 3D depth information that placed two avatars at a constant distance of two virtual meters in front of the observer in depth and were located one virtual meter to the left and right of the observer’s straight-ahead. The two avatars could both be the “self” or “other” avatar, or one of each, with either 1:1 or decoupled motion applied to them. Eight conditions made up from these arrangements were each repeated ten times using the method of constant stimuli for a total of 80 trials (Table in Figure 4). Participants were encouraged by the experimenter to move around and interact with the avatars for ten seconds after which instructions on the screen prompted the participants to indicate whether the avatar to their left or their right most represented their self using their corresponding left or right hand with an “ok” gesture that was recorded and coded as a left or right judgement. Following the experiment we asked participants to report their cybersickness level that were collected using a quick verbal report (Fast Motion Sickness scale, FMS; Keshavaraz & Hecht, 2011). FMS scores are measured as a verbal response to the following question: “On a scale from 0-20 with 0 being no sickness and 20 being severe sickness, how do you feel?”

### Analysis

Probabilities of selecting one avatar type versus another were averaged over 20 trials per condition (%self normal vs. %self decoupled; %other normal vs. other decoupled; %self normal vs. other decoupled; %self decoupled vs. other normal). As the dataset failed Shapiro-Wilk tests for normality and Bayesian statistics can handle outliers by describing the data as heavy tailed distributions instead of normal distributions (Kruschke 2013), we used Bayesian one-sample t-tests to assess whether probabilities of avatar preference for each avatar pair were significantly different from chance using JASP v0.8.0.1. Here Bayes Factors (BF; default Cauchy prior width = 0.707) provide a numerical value that quantifies how well a hypothesis (*H*_1_; preferred avatar significantly different from chance) predicts the data relative to a competing null hypothesis (*H*_0_; avatar preference at chance), where a BF_10_ between 0 and 1, indicates support for the *H*_0_, and a BF_10_ greater than 1 indicates support for the *H*_1_. A two (motion: normal vs. decoupled) by two (avatar type: self vs. other) Bayesian repeated measures ANOVA was used to assess for main effects and interactions.

## Results

All participants were able to understand and complete the task. Reported cybersickness was low (Average FMS: 1.625, SD = 1.41). As shown in Figure 5, when participants were confronted with two self-avatars, avatar preference was in the direction of the self-avatar with normal motion (mean: 54.5%, s.e. 5.1) but at chance (support for *H*_1_; BF_10_ = 0.446). When confronted with two other-avatars, participants preferred the avatar with normal 1:1 motion (mean: 60%, s.e. 3.8; support for *H*_1_; BF_10_ = 2.643). As expected participants preferred the self-avatar when participants were confronted with their self-avatar and the other-avatar, with the other-avatar having decoupled motion (mean: 76%, s.e. 7.8; support for *H*_1_; BF_10_ = 5.973).

**Figure 5:**
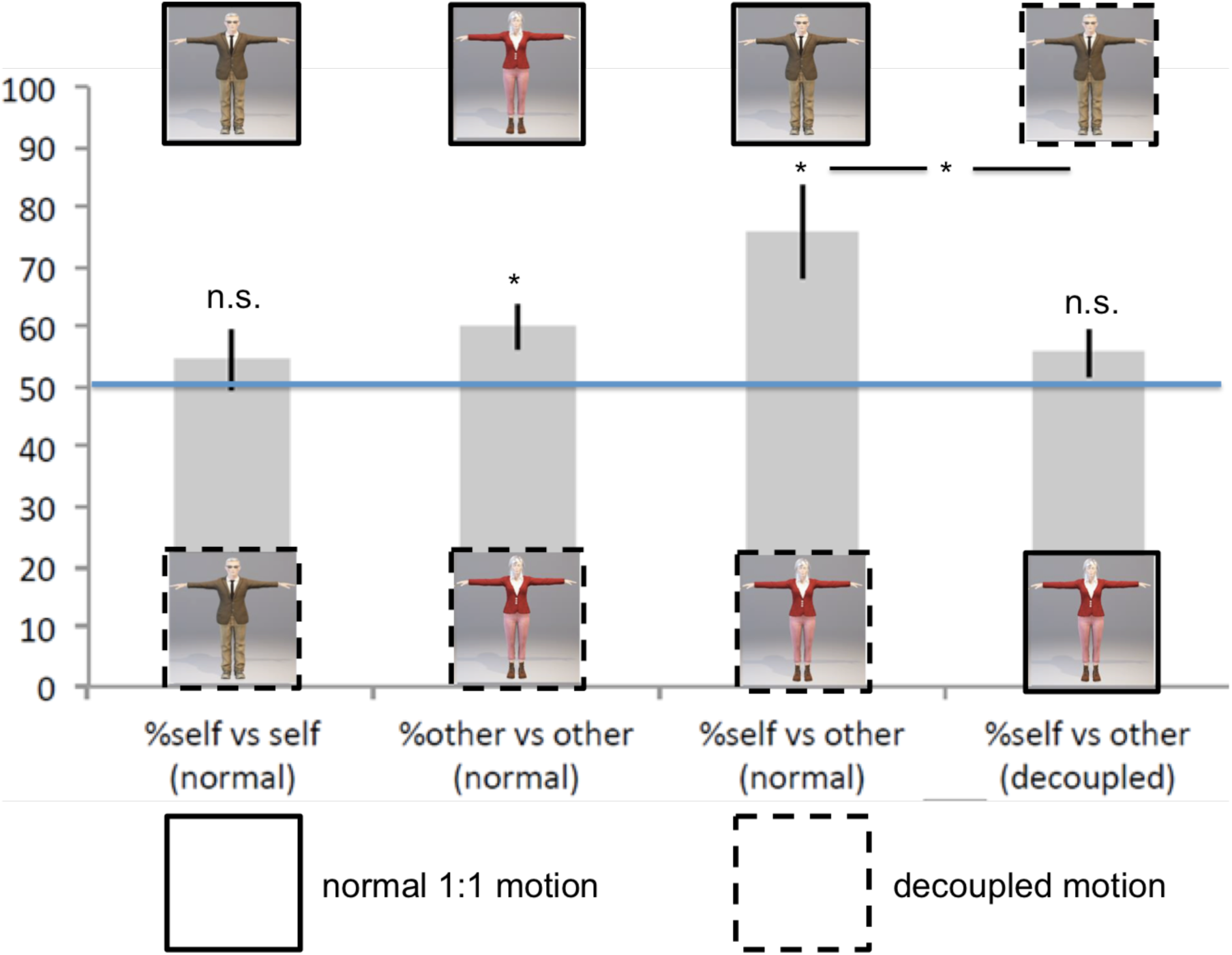
Experimental results. Probability of selecting the “self” avatar with 1:1 motion compared to the self avatar with decoupled motion was not different from chance (far left). The “other” avatar with 1:1 motion was significantly preferred over the other avatar with decoupled motion (middle left). The self avatar with 1:1 motion was strongly preferred to the other avatar with decoupled motion (middle right), however, this preference returned to chance probability when decoupled motion was applied to the self avatar and 1:1 motion to the other avatar indicating an interaction between self-motion and sex identification for the sense of self.

Surprisingly, when participants were confronted with their self-avatar and the other-avatar, with the other-avatar having normal motion, participants no longer preferred their self-avatar by returning to chance preference (mean: 55.7%, s.e. 4; support for *H*_1_; BF_10_ = 0.687). While there was no main effect of motion (support for *H*_1_; BF_10_ = 0.769), there was a main effect of avatar type (support for *H*_1_; BF_10_ = 2.862), as well as a significant interaction between motion and avatar type (support for *H*_1_; BF_10_ = 2.53), driven by a significant difference between the two mixed avatar conditions (Bayesian paired sample t-test: support for *H*_1_; BF_10_ = 1.547).

## Discussion

When participants in our virtual reality experiment encountered two avatars that mirrored their actions, participants were more likely to select the avatar that they created to resemble themselves and that moved in close correspondence with their own self-motion. By decoupling visual self-motion feedback and by changing the avatar’s apparent biological sex characteristics, however, our results from all conditions show that visual self-motion feedback and biological sex identification affect the sense of self in virtual reality through a significant interaction. This was most compelling where participants were equally likely to identify with either the same sex or opposite sex avatar when visual self-motion feedback was slightly decoupled for the same sex avatar. Together, these findings are consistent with previous research in healthy individuals showing that the sense of self and embodiment are relatively plastic and can lead to compelling perceptions of having ownership over additional limbs (Guterstam et al., 2011), body parts (Tsakiris et al., 2010), and whole bodies, even those of the opposite sex (Petkova and Ehrsson, 2008).

Why is the sense of self so malleable? The central nervous system is challenged with constantly having to detect signals from the environment and from within the body to form a coherent representation of the self in its surroundings (Ernst and Bülthoff, 2004). As these signals are detected from different sensory organs, our perceptual systems are continuously integrating and interpreting multisensory information to compute the spatial relationship between our body and the world around us (Ehrsson, 2010). This process could be quite computationally demanding and potentially create large delays in signal processing and action planning, which would be dangerous to the survival of any organism. However, the central nervous system can instead rely on internal models and prior knowledge about the body and the environment at the cost of misrepresenting the self. Such a framework can explain illusions that involve displacement (Botvinick and Cohen, 1998; Ehrsson, 2007), changes in size (Linkenauger et al., 2010), and experiences of being outside of one’s body (Ehrsson, 2007; Lopez et al., 2008; Blanke, 2012), provided there are no gross violations concerning the basic structure of the human body (Tsakiris et al., 2010; Guterstam et al., 2011).

While the study presented here has bearing on both the philosophical and neuroscientific perspectives on the sense of self, there are some limitations that should be noted. First, this experiment was put together as part of a workshop for a visiting Korean group of students. While every effort was made to fully translate all instructions and ensure participants fully understood the nature of the experiment it is possible that the circumstances of this data collection could have some bearing on the results. While the sample size was also small due to the convenience sample of the visiting group, we used a robust Bayesian statistical framework that is capable of equally assessing both the null and alternative hypotheses. We do suggest that future research should replicate this experiment with a larger sample size under different recruitment circumstances. More importantly, we did not collect measures of presence nor did we collect gender preferences from our participants. This we also suggest should be addressed in future experimentation as our unique virtual reality paradigm would be ideally suited to assess to what degree to participants identify their self with their same or opposite sex. Moreover, the paradigm could also enable measurement of the sense of self for changes of other characteristics related to body size, satisfaction, and even adopting non-human avatars that could be embodied. Indeed parametric evaluation of the extend to which visual self-motion is decoupled from true motion in addition to changes in the trait characteristics of avatars would allow for the construction of predictive computational models for embodiment in virtual reality.

## Acknowledgments

We are grateful to Daniel Cheriyan for helping to conduct this experiment as well as Hyun Suk Choi, Jaeyoung Kang, and Yoolim Lee who provided English-Korean translation services in preparation for and during the experiment. This research was supported by a research grant to MBC from the National Engineering Research Council of Canada (#RGPIN-05435-2014), and a research grant to JT from the Social Sciences and Humanities Research Council of Canada, and the Canada Research Chairs program.

## References

Aglioti, S. M. & Candidi, M. (2011). Out-of-place bodies, out-of-body selves. Neuron, 70, 173–175 https://doi.org/10.1016/j.neuron.2011.04.006

Blanke, O. (2012). Multisensory brain mechanisms of bodily self-consciousness. Nature Reviews Neuroscience, 13, 556–571 https://doi.org/10.1038/nrn3292

Blanke, O. & Metzinger, T. (2009). Full-body illusions and minimal phenomenal selfhood. Trends in Cognitive Sciences, 13, 7–13 https://doi.org/10.1016/j.tics.2008.10.003

Blanke, O., Slater, M. & Serino, A. (2015). Behavioral, neural, and computational principles of bodily self-consciousness. Neuron, 88:, 145–166 https://doi.org/10.1016/j.neuron.2015.09.029

Botvinick, M. & Cohen, J. (1998). Rubber hands ‘feel’ touch that eyes see. Nature 391, 756 http://dx.doi.org/10.1038/35784

Ehrsson, H.H. (2007). The experimental induction of out-of-body experiences. Science, 317, 1048 https://doi.org/10.1126/science.1142175

Ehrsson, H.H. (2010). The Concept of Body Ownership and its Relation to Multisensory Integration. In: MIT Handbook of Multisensory Integration. Boston, USA: MIT Press ISBN: 9780262033213

Ehrsson, H.H., Holmes, N.P. & Passingham, R.E. (2005). Touching a rubber hand: feeling of body ownership is associated with activity in multisensory brain areas. Journal of Neuroscience, 25, 10564–10573 https://doi.org/10.1523/JNEUROSCI.0800-05.2005

Ernst, M.O. & Bülthoff, H.H. (2004). Merging the senses into a robust percept. Trends in Cognitive Sciences, 8, 162–169 https://doi.org/10.1016/j.tics.2004.02.002

Ferrè, E.R. & Haggard, P. (2015). Vestibular–Somatosensory Interactions: A Mechanism in Search of a Function? Multisensory Research, 28, 559–579 https://doi.org/10.1163/22134808-00002487

Gallagher, S. (2000). Philosophical conceptions of the self: implications for cognitive science. Trends in Cognitive Sciences, 4, 14–21 https://doi.org/10.1016/S1364-6613(99)01417-5

Gallup, G.G. (1970). Chimpanzees: self-recognition. Science, 167, 86–87 https://psycnet.apa.org/doi/10.1126/science.167.3914.86

Guterstam, A., Petkova, V.I. & Ehrsson, H.H. (2011). The illusion of owning a third arm. PLoS One, 6, e17208 https://doi.org/10.1371/journal.pone.0017208

Hart, J.W. & Scassellati, B. (2012). Mirror perspective-taking with a humanoid robot. In: Twenty-Sixth AAAI Conference on Artificial Intelligence.

Kaliuzhna, M., Vibert, D., Grivaz, P. & Blanke, O. (2015). Out-of-body experiences and other complex dissociation experiences in a patient with unilateral peripheral vestibular damage and deficient multisensory integration. Multisensory Research, 28, 613–635 https://doi.org/10.1163/9789004342248_013

Kruschke, J.K. (2013). Bayesian estimation supersedes the t test. Journal of Experimental Psychology: General, 142, 573–603 https://psycnet.apa.org/doi/10.1037/a0029146

Lim, S. & Reeves, B. (2009). Being in the game: effects of avatar choice and point of view on psychophysiological responses during play. Media Psychology, 12, 348–370 https://doi.org/10.1080/15213260903287242

Linkenauger, S.A., Ramenzoni, V. & Proffitt, D.R. (2010). Illusory shrinkage and growth: body-based rescaling affects the perception of size. Psychological Science, 21, 1318–1325 https://doi.org/10.1177%2F0956797610380700

Longo, M.R., Cardozo, S. & Haggard, P. (2008). Visual enhancement of touch and the bodily self. Consciousness and Cognition, 17, 1181–1191 https://doi.org/10.1016/j.concog.2008.01.001

Lopez, C. (2015). Making sense of the body: The role of vestibular signals. Multisensory Research, 28, 525–557 https://doi.org/10.1163/22134808-00002490

Maravita, A., Iriki, A. (2004). Tools for the body (schema). Trends in Cognitive Sciences, 8, 79–86 https://psycnet.apa.org/doi/10.1016/j.tics.2003.12.008

Maister, L., Slater, M., Sanchez-Vives, M. V., & Tsakiris, M. (2015). Changing bodies changes minds: owning another body affects social cognition. Trends in Cognitive Sciences, 19, 6–12 https://doi.org/10.1016/j.tics.2014.11.001

Mölbert, S.C., Thaler, A., Mohler, B.J., Streuber, S., Romero, J., Black, M.J., Zipfel, S. Karnath, H-O. & Giel K.E. (2018). Assessing body image in anorexia nervosa using biometric self-avatars in virtual reality: Attitudinal components rather than visual body size estimation are distorted. Psychological Medicine, 48, 642–653 https://doi.org/10.1017/S0033291717002008

Petkova, V.I. & Ehrsson, H.H. (2008). If I were you: perceptual illusion of body swapping. PloS One, 3, e3832 https://doi.org/10.1371/journal.pone.0003832

Reiss, D. & Marino, L. (2001). Mirror self-recognition in the bottlenose dolphin: a case of cognitive convergence. Proceedings of the National Academy of Sciences, 98, 5937–5942 https://doi.org/10.1073/pnas.101086398

Sanchez-Vives, M.V. & Slater, M. (2005). From presence to consciousness through virtual reality. Nature Reviews Neuroscience, 6, 332–339 https://doi.org/10.1038/nrn1651

Suzuki, K., Garfinkel, S. N., Critchley, H. D., & Seth, A. K. (2013). Multisensory integration across exteroceptive and interoceptive domains modulates self-experience in the rubber-hand illusion. Neuropsychologia, 51, 2909–2917 https://doi.org/10.1016/j.neuropsychologia.2013.08.014

Synofzik, M., Vosgerau, G. & Newen, A. (2008). I move, therefore I am: a new theoretical framework to investigate agency and ownership. Consciousness and Cognition, 17, 411–424 https://doi.org/10.1016/j.concog.2008.03.008

Tsakiris, M., Carpenter, L., James, D. & Fotopoulou, A. (2010). Hands only illusion: multisensory integration elicits sense of ownership for body parts but not for noncorporeal objects. Experimental Brain Research, 204, 343–352 https://doi.org/10.1007/s00221-009-2039-3

Tsakiris, M. & Haggard, P. (2005) The rubber hand illusion revisited: visuotactile integration and self-attribution. Journal of Experimental Psychology: Human Perceptaion and Performance, 31, 80–91 https://psycnet.apa.org/doi/10.1037/0096-1523.31.1.80

